# Complex admixture preceded and followed the extinction of wisent in the wild

**DOI:** 10.1101/059527

**Authors:** Karolina Węcek, Stefanie Hartmann, Johanna L. A. Paijmans, Ulrike Taron, Georgios Xenikoudakis, James A. Cahill, Peter D. Heintzman, Beth Shapiro, Gennady Baryshnikov, Aleksei N. Bunevich, Jennifer J. Crees, Roland Dobosz, Ninna Manaserian, Henryk Okarma, Małgorzata Tokarska, Samuel T. Turvey, Jan M. Wójcik, Waldemar Żyła, Jacek M. Szymura, Michael Hofreiter, Axel Barlow

## Abstract

Retracing complex population processes that precede extreme bottlenecks may be impossible using data from living individuals. The wisent (*Bison bonasus*), Europe’s largest terrestrial mammal, exemplifies such a population history, having gone extinct in the wild but subsequently restored by captive breeding efforts. Using low coverage genomic data from modern and historical individuals, we investigate population processes occurring before and after this extinction. Analysis of aligned genomes supports the division of wisent into two previously recognised subspecies, but almost half of the genomic alignment contradicts this population history as a result of incomplete lineage sorting and admixture. Admixture between subspecies populations occurred prior to extinction and subsequently during the captive breeding program. Admixture with the *Bos* cattle lineage is also widespread but results from ancient events rather than recent hybridisation with domestics. Our study demonstrates the huge potential of historical genomes for both studying evolutionary histories and for guiding conservation strategies.

## Introduction

The last known wild wisent, or European bison (*Bison bonasus*), was shot and killed in 1927, marking the extinction of this species in the wild (Pucek, 1991). As a result of an intensive captive breeding program and a series of re-establishments, today the species again occupies part of its former range in Central and Eastern Europe. The total population of free-ranging wisent now stands at 5,553 (Raczyński, 2014), and the International Union for Conservation of Nature no longer considers the wisent as an endangered species (IUCN 2008).

The decline of the original wisent population was protracted, and their subsequent restitution has been complex. In historical times, wisent ranged extensively across semi-open habitats and broadleaved, mixed and coniferous forests in Western Europe, from what is today France in the west to the Volga River and the Caucasus in the east, with the northernmost range limits around 60° north (Fig. 1; Kuemmerle et al., 2011; Kerley et al., 2012; Bocherens et al., 2015). However, ongoing habitat fragmentation and overhunting eradicated most populations. By the end of the 19th century, there were only two populations of wisent left in the wild that were assigned to separate subspecies: in Białowieża Forest (Lowland wisent, *B. b. bonasus*) and in the western Caucasus Mountains (Caucasian wisent, *B. b. caucasicus*). Finally, even these populations collapsed; the last wild Lowland wisent was shot in Poland in 1919 followed by the last Caucasian animal in 1927 (Pucek, 1991).

**Figure 1.**
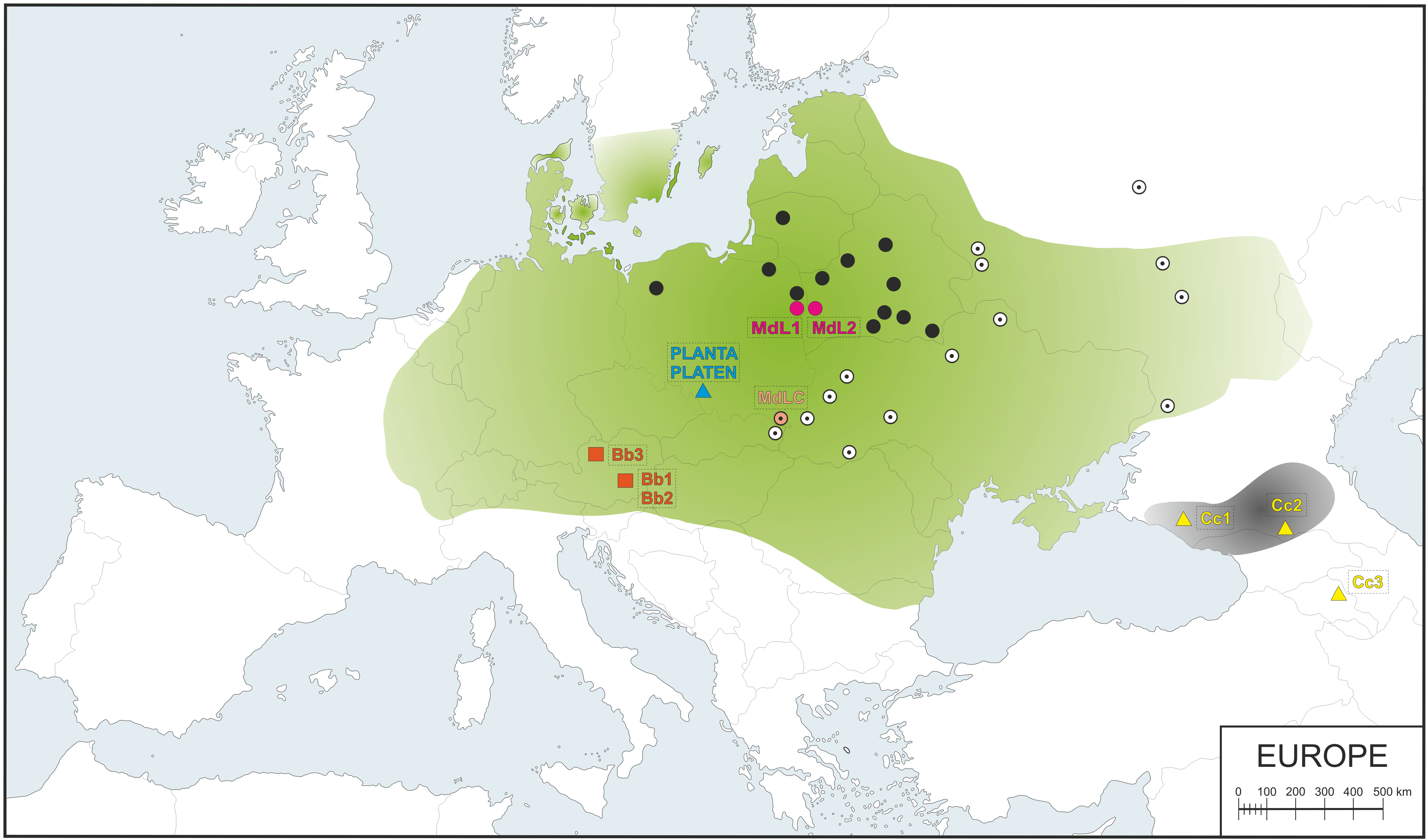
Map of Western Europe showing the putative historical range of Lowland wisent (shaded green) and Caucasian wisent (shaded grey) based on bone remains and written records (according to Benecke et al., 2005; Kuemmerle et al., 2011; Tokarska et al., 2011; Bocherens et al., 2015) and sample locations. Black circles indicate contemporary free-ranging modern L line herds and white circles indicate modern LC line herds. Purple and peach circles denote the locations of investigated modern L (MdL1, MdL2) and LC (MdLC) line wisent, respectively, orange squares show the location of the Holocene wisent (Bb1-3) and blue and yellow triangles indicate historical founding wisent from the Pszczyna population (PLANTA and PLATEN) and the extinct Caucasian wisent (Cc1-3), respectively.

In 1924, the captive population consisted of only 54 individuals (29 males and 25 females). However, detailed analysis of pedigrees (Olech 2009, Slatis 1960) has shown that some individuals in this population were the direct descendents of others, and that some individuals left no modern descendents. Thus, the true founding population of wisent was considerably smaller, and is thought to comprise of just 12 individuals (Slatis 1960). All but one of these 12 founders were Lowland wisent, almost half of which came from a population established in 1865 in Pless (now Pszczyna, Poland). The remaining founder was a Caucasian wisent bull named M100 KAUKASUS, that represented the last surviving pure Caucasian wisent in captivity. The modern herds that are derived from this founding population are managed as two separate genetic lines. The Lowland line (L) derives from 7 Lowland founders (4 males and 3 females), and is thus considered to represent a pure Lowland wisent lineage. The Lowland-Caucasian (LC) line originates from all 12 founders (5 males, 7 females), which included the last remaining Caucasian wisent bull (Slatis, 1960). Descendants of the LC line thus represent a mixture of Lowland and Caucasian wisent ancestry.

Although the wisent restitution undoubtedly represents a tremendous conservation success, several factors may limit the long-term viability of the species, many of which are applicable to ex-situ conservation strategies in general. A factor that has received particular attention is that of reduced genetic variability, which may be correlated with a lowered resistance to disease and parasites in wisent (e.g.: Krasińka & Krasińki, 2007 and references therein; Adaszek et al., 2014; Karbowiak et al., 2014a; 2014b and the references therein; Krzysiak et al., 2014; Majewska et al., 2014; Moskwa et al., 2014; Oleński et al., 2015; Panasiewicz et al., 2015), and also seriously impacts conservation programs for other threatened species (Altizer et al., 2007). Although genetic variability and geneflow among living wisent herds has been investigated using limited sets of genetic markers (e.g. Gralak et al., 2004; Luenser et al., 2005; Radwan et al., 2007; Wójcik et al., 2009; Babik et al., 2012; Tokarska et al., 2009a, 2009b, 2015), to date no published study has investigated the genetic composition of extant wisent at the level of the complete genome. Similarly, while preserved specimens of some founding individuals that may retain genetic information still exist within museum collections, the genetic variability among these founders and their subsequent contribution to living herds remains unquantified.

A second threat for wisent is potential hybridisation with domestic cattle (*Bos taurus*). Although F1 hybrid bulls are sterile, hybrid female offspring are not (Basrur, 1968), and would therefore have the potential to reintegrate back into the wild population. It is known that domestic cattle were grazed in the Białowieża Forest of Poland and Belarus for many years at the same time as wisent were present there, though no births of hybrids in natural conditions were recorded (Krasińka & Krasińki 2007). Hybridisation with domestics in general is of wider concern for threatened species (Wolf et al., 2001). Well known examples of this kind of hybridisation include Przewalski’s horse and domestic horse (Sarkissian et al., 2015), mouflon and domestic sheep (Lühken et al., 2009), wolf/coyote and dog (Bohling & Waits, 2011), wildcat and domestic cat (Steyer et al., 2016), and game birds and domestic fowl (Arrieta et al., 2013). However, in the case of wisent, the extent of cattle admixture into living herds remains undetermined, as does the admixture that may have occurred prior to the establishment of the captive breeding program. Such information is critical to assess the magnitude of the threat that cattle admixture represents to the long term viability and integrity of the species.

Here we present low-coverage whole genome sequencing data from both modern genetic lines, as well as from four historical samples representing two of the original founding individuals and two individuals of the now extinct Caucasian wisent subspecies. Using these data, we compare the relative contributions of the founding individuals to the genomes of living wisent. We also determine the magnitude and distribution of Caucasian wisent ancestry in modern and founding individuals. Furthermore, we uncover evidence of admixture with the cattle/aurochs lineage that has resulted in a significant component of mixed ancestry in the genomes of both founding and modern wisent. Our results have important implications for both understanding the evolutionary history and for the future conservation management of wisent. Finally, we demonstrate the huge potential of genomic approaches, in particular applied to historical samples, for conservation management of endangered species.

## RESULTS

### Sequencing of wisent genomes

We conducted shotgun sequencing of wisent genomes using Illumina technology and mapped the resulting sequence reads to a reference genome assembly of the Asian water buffalo, *Bubalus bubalis* (GenBank accession no. GCA_000471725.1), which represents an outgroup to the wisent/cattle clade (Fig. S1; Hassanin et al., 2013). In total we sequenced seven individuals that we divide into three categories. Detailed sample information, including provenance, is provided in Table 1 and sample localities are shown in Figure 1.

**Table 1.**
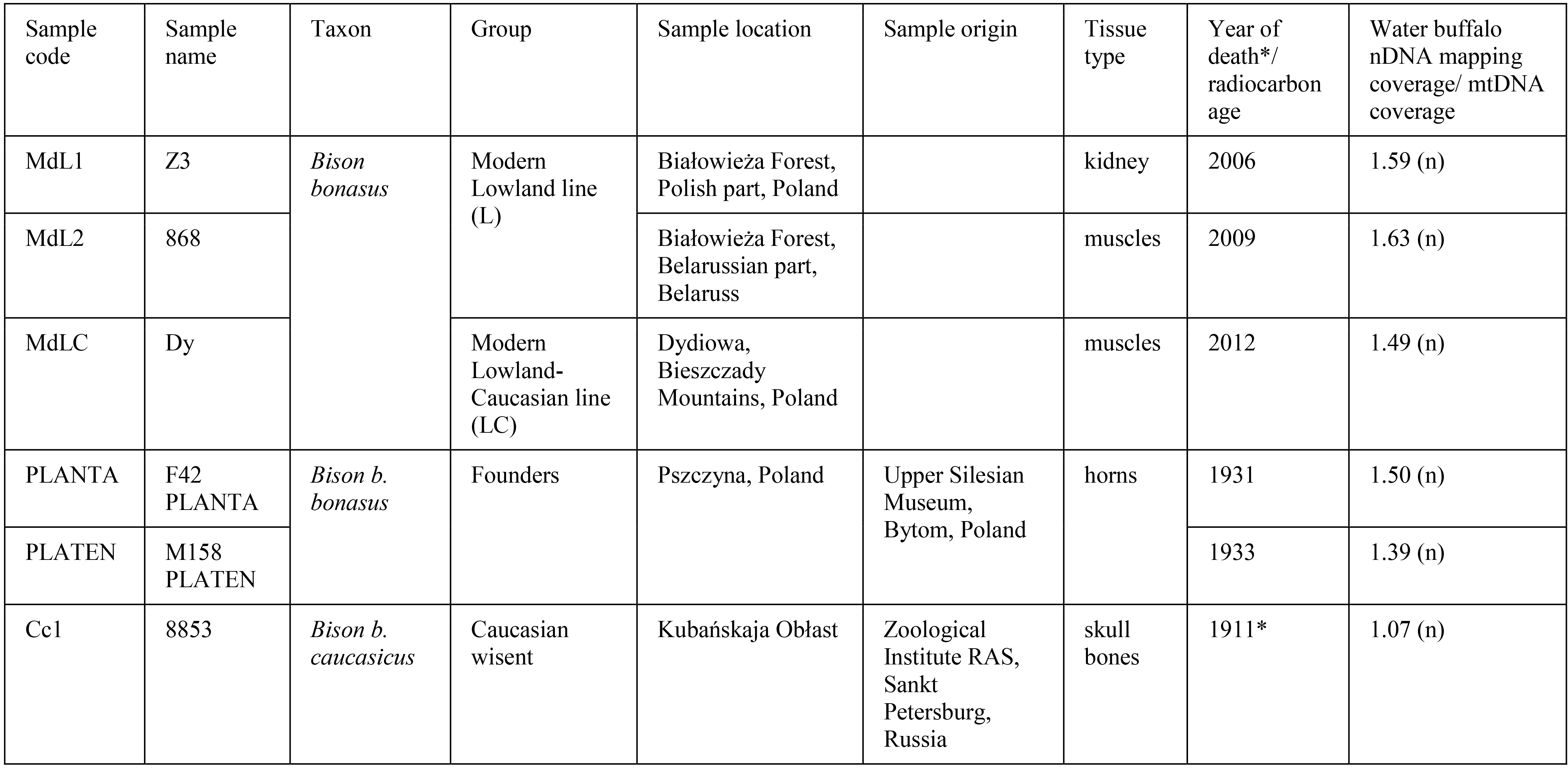
Basic information about sampled individuals. The last column is the average coverage for each sample after mapping it to either the water buffalo (*Bubalus bubalis*) nuclear reference genome (GenBank accession no. GCA_000471725.1) or the American bison (*Bison bison*) mitochondrial reference genome (GenBank accession no. NC_012346.1).

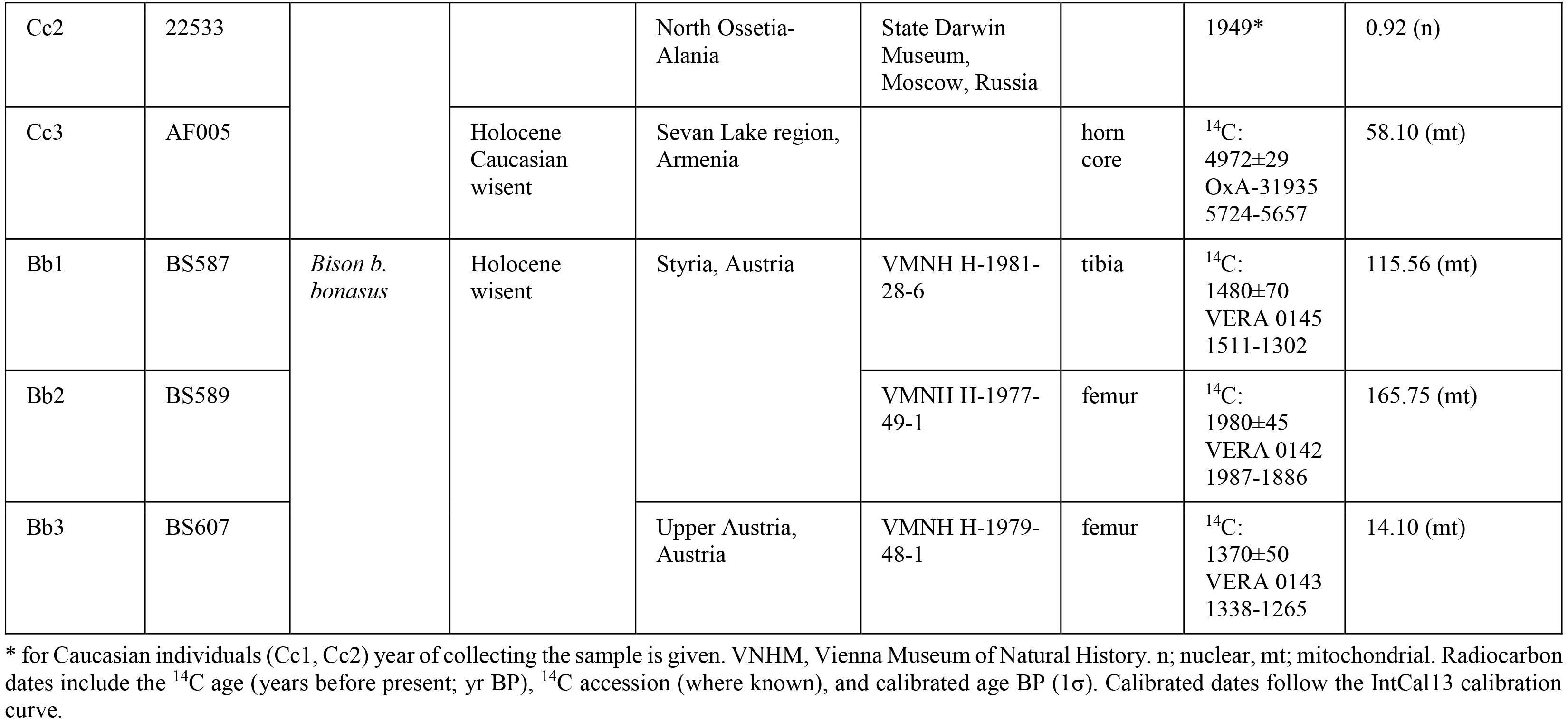

*Modern wisent* – three modern individuals representing both genetic lines. These comprise two individuals from the L line (MdL1, mean read depth 1.59x; and MdL2, mean read depth 1.63x; from the Polish and Belarussian parts of the Białowieża Forest, respectively) and one individual from the LC line (MdLC, from Dydiowa in the Bieszczady Mountains, mean read depth 1.49x).

*Founding wisent* – two individuals assignable to the Lowland wisent subspecies *B. b. bonasus* from the initial breeding population originating from Pszczyna, both of which contributed to the establishment of both the L and the LC genetic lines: foundress F42 PLANTA (1904-1931, mean read depth 1.82x) and her male offspring, M158 PLATEN (1926-1933, mean read depth 1.36x), who was fathered by another founder M45 PLEBEJER (1917-1937).

*Caucasian wisent* – two individuals from the early 1900’s representing the now extinct Caucasian wisent subspecies *B. b. caucasicus* (Cc1, from Kubańskaja Obłast, mean read depth 1.17x; and Cc2, from North Ossetia-Alania, mean read depth 0.92x).

For each individual, we collapsed mapped reads into a single pseudo-haploid genome sequence by randomly selecting a single high quality nucleotide from the read stack at each position of the reference genome, following the procedure described by Cahill et al. (2013; 2015). This procedure disregards heterozygous positions, where only one allele will be sampled, but should not introduce any biases in allele sampling. Ancient DNA fragments frequently contain miscoding lesions resulting from postmortem DNA degradation, the most common of which involves the deamination of cytosine to uracil, which causes C to T substitutions in the resulting data (e.g. Dabney et al., 2013b). This pattern is present at varying levels in sequence data from our historical samples (Fig. S2-S5), and so we restricted all subsequent analyses to transversion sites only to avoid any confounding effects of DNA damage.

### Genomic divergence

We investigated patterns of nuclear genomic divergence among wisent by conducting pairwise comparisons of the number of transversion differences occurring along a sliding window of 1Mb, producing a distribution of genomic divergence for each wisent pair. The resulting probability densities showed that nuclear genomic divergence is broadly similar among all modern and founding wisent (Fig. 2a). The two founding individuals, PLANTA and PLATEN, are somewhat less diverged from one another than either is from all modern wisent (Fig. 2a), reflecting their mother-son relationship. Slightly increased divergence is observed between the modern LC line individual (MdLC) and all modern and founding wisent (Fig. 2a), which may reflect the increased component of Caucasian wisent ancestry in this individual resulting from the captive breeding program.

**Figure 2.**
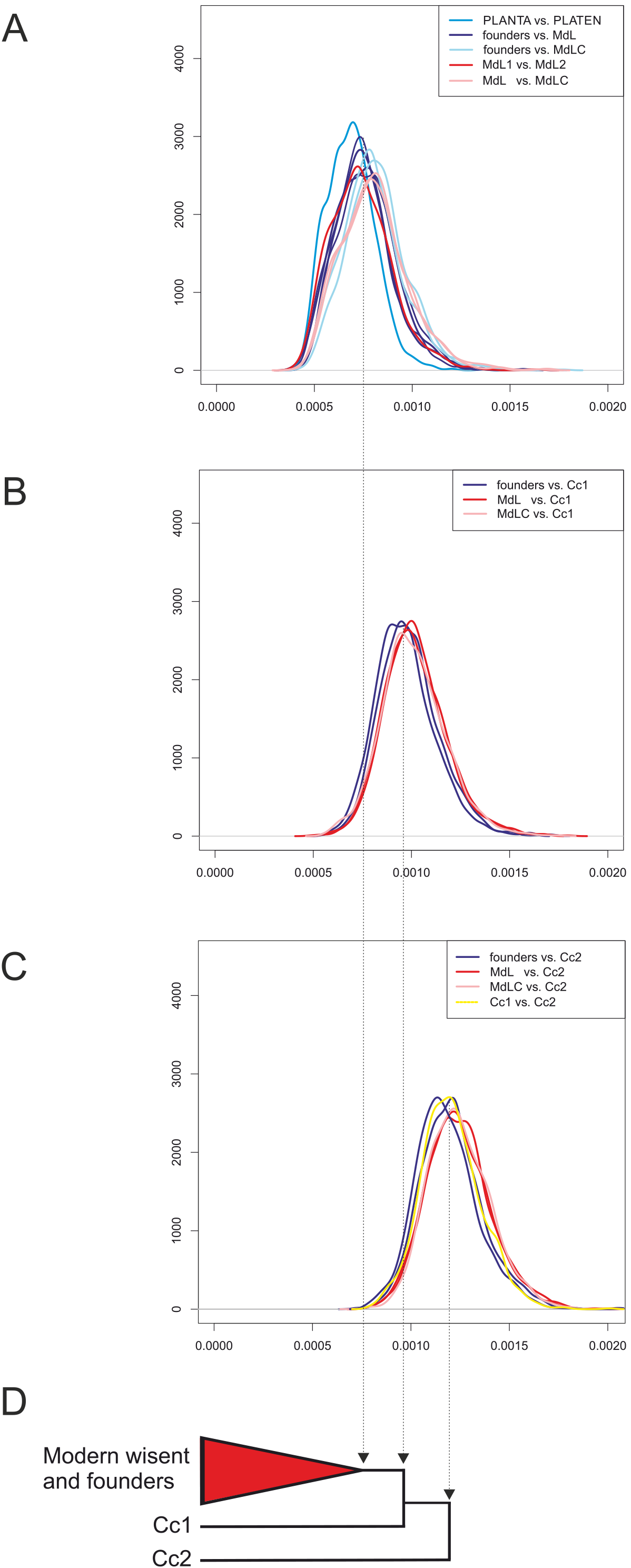
Pairwise genomic divergence among wisent. A, B, C, show probability densities for pairwise transversion divergence (x axes) along a sliding window of 1Mb. Individual plots show all pairwise comparisons among modern and founding individuals (A); comparisons of modern and founding individuals and Caucasian wisent Cc1 (B); and comparisons of Caucasian wisent Cc2 and all other individuals (C). Specific comparisons discussed in the text are identified by colours, according to the key at the top right of each plot. Schematic neighbour-joining phylogeny based on whole genome distances (D).

Genomic divergence between Caucasian and both modern and founding wisent greatly exceeds that occurring between the latter two groups (Fig. 2a-c). Substantial divergence is also found between the two Caucasian wisent individuals. One of these Caucasian wisent (Cc1) was found to be less diverged from modern and founding wisent than other Caucasian wisent individual (Cc2), suggesting the presence of not only substantial genetic diversity but also substantial population structure in the extinct Caucasian wisent subspecies.

We also investigated mitochondrial genome variability among all individuals subjected to nuclear genome sequencing, in addition to eight other modern wisent (Tab. S1). Sequence analysis revealed that all investigated modern wisent, both founding wisent, and a single historical Caucasian wisent (Cc1), share a single haplotype. The haplotype occurring in the second historical Caucasian wisent (Cc2) differed from this widely shared haplotype by a single transition site. These results hint at a major loss of mitochondrial haplotype diversity prior to the extinction of wisent in the wild. This inference is supported by additional haplotypes that we recovered from three ancient middle Holocene wisent from Austria and one ancient middle Holocene Caucasian wisent from Armenia (Tab. 1). All these ancient haplotypes are substantially divergent from those found in modern and historical wisent, suggesting a substantial loss of haplotype diversity, potentially within the last ~1500 years.

Neighbour-joining phylogenetic analysis of total nuclear genomic divergence supports paraphyly of Caucasian wisent (Fig. 2d), as does analysis of mitochondrial haplotypes (Fig. S1). We further investigated the population history of wisent by dividing aligned nuclear genome sequences into non-overlapping 1MB blocks and subjecting each block to maximum-likelihood phylogenetic analysis. For this analysis, we included both Caucasian wisent and both modern L line wisent as representatives of the Lowland wisent subspecies, with water buffalo as outgroup. Founding wisent and the modern LC line wisent were not included to avoid any confounding effects of direct ancestor-descendent relationships and documented Caucasian wisent introgression (Slatis, 1960) on phylogenetic interpretation, respectively. We found that 57% of the investigated genomic blocks support reciprocal monophyly of Caucasian and Lowland wisent (Fig. 3). We therefore conclude that this most likely represents the true population history. All alternative topologies occur, individually, at a much lower frequency. Nevertheless, almost half of the genome sequence alignment of these individuals contradicts the true population history.

**Figure 3.**
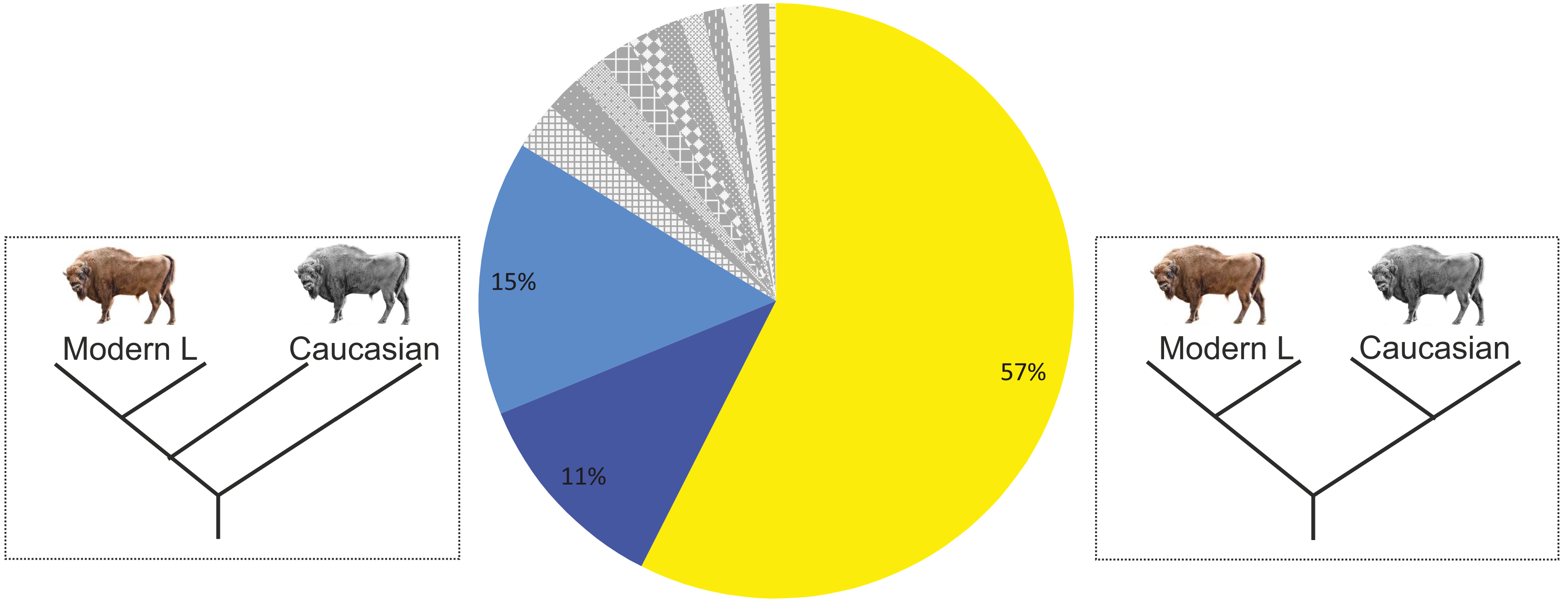
Population history of Lowland and Caucasian wisent, estimated using two representatives of each. The pie chart shows the percentage of 1Mb genomic blocks supporting each alternatively rooted tree topology. The indicates fraction of genome blocks returning both wisent subspecies as reciprocally monophyletic. Dark and light blue colours show the next most frequently encountered topologies in which Caucasian wisent are paraphyletic (dark blue: Cc2 most divergent, and light blue: Cc1 most divergent, the first of which is compatible with estimates pairwise genomic divergence (see Fig 2d).

In order to interpret wisent genomic divergence in the context of total species genetic diversity, we obtained data from the NCBI Short Read Archive for seven domestic cattle and seven yak (*Bos grunniens*; Tab. S4) and subjected them to the same analysis pipeline. Genomic divergence among modern wisent was found to be similar to that found among domestic cattle, and exceeded that found among yak (Fig. 4). We conducted equivalent comparisons for modern wisent and these other bovid species but with the inclusion of transition as well as transversion sites. Interestingly, the distribution of genomic divergence for pairs of wisent was bimodal in all cases (Fig. 4), but the relative levels of genomic divergence between species were similar to that measured using only transversion sites.

**Figure 4.**
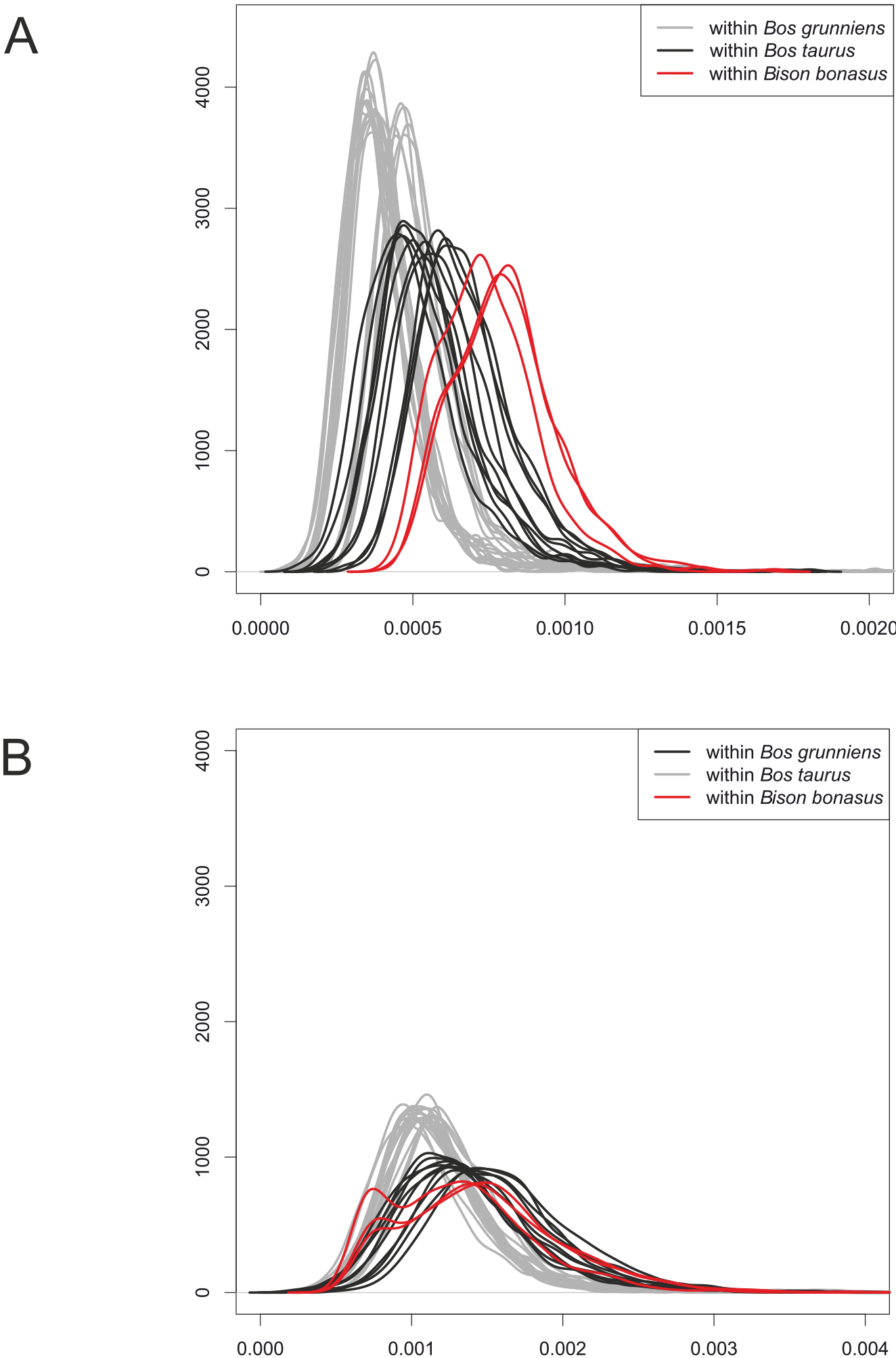
Comparison of pairwise genomic divergence within three bovid species: wisent (*Bison bonasus*), domestic cattle (*Bos taurus*), and yak (*Bos grunniens*). Probability densities were calculated along a sliding window of 1Mb from transversions only (A), and from transitions and transversions (B). For wisent, only modern samples included.

### Wisent geneflow and admixture

We investigated patterns of admixture among wisent using the *D* statistic test for admixture (Green et al., 2010). This test identifies any imbalance in the number of derived alleles that either of two closely related individuals share with a candidate introgressor. A significant excess of derived alleles shared between one individual and the introgressor provides evidence of admixture between them (Durand et al., 2011). For all *D* statistic tests, we used water buffalo (*Bubalus bubalis*) as outgroup for allele polarisation.

We first investigated patterns of derived allele sharing among modern wisent, and found no statistically significant signal of admixture between the modern LC line individual and either modern L line individual (Tab. S5). Between modern and founding wisent, we found that modern L line wisent share a significantly greater number of derived alleles with founding wisent than the modern LC line individual does (Fig. 5a), indicating a greater contribution of the two founding wisent investigated here to the L line, relative to the LC line, which is consistent with pedigrees (Slatis, 1960).

**Figure 5.**
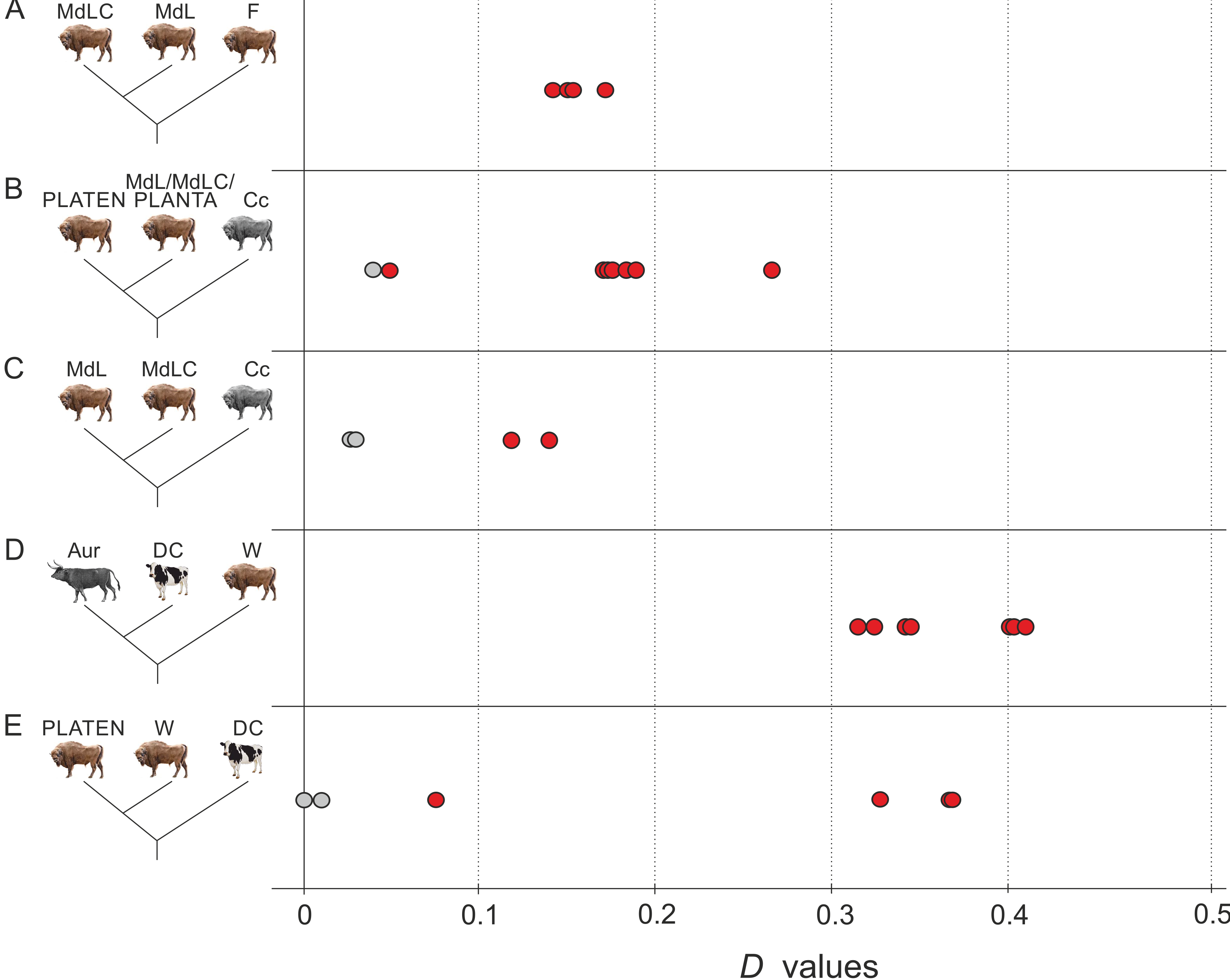
Results of *D* statistic analysis. Red and grey points show significant and nonsignificant *D* values (x axis), respectively, and show: the genetic contribution of the founders (F) to the modern individuals (A); Caucasian wisent (Cc) admixture with modern L and LC herds (MdL and MdLC) and one founder, PLANTA, relative to the least Caucasian admixed wisent, founder PLATEN (B); Caucasian wisent admixture among modern wisent (C); apparent cattle (DC) admixture with all investigated wisent (W) relative to aurochs (Aur) (D); variance in cattle/aurochs admixture among wisent (W) compared to PLATEN (E). Detailed *D* statistic results are provided in Supporting Table S5.

We then investigated admixture involving Caucasian wisent. We found a significant excess of derived allele sharing between one founding wisent, PLANTA, and one Caucasian wisent, Cc2, relative to the other founding wisent, her son PLATEN (Fig. 5b). This indicates that a proportion of the genome of PLANTA can be attributed to admixture with Caucasian wisent. Furthermore, we can deduce that PLEBEJER, the father of PLATEN, must have possessed a lower level of Caucasian wisent admixture than PLANTA, and that PLATEN himself was likely admixed to some degree through inheritance from PLANTA. The detection of admixture involving one Caucasian wisent (Cc2) but not the other (Cc1) further supports the existence of genetic structure in Caucasian wisent inferred from estimates of genomic divergence (Figs. 2b & 2c).

Next, we investigated evidence of Caucasian wisent admixture among modern wisent. Consistent with expectations, we found that the modern LC line individual (MdLC) shares an excess of derived alleles with one of the Caucasian wisent (Cc1) relative to modern L line individuals (Fig. 5c). We did not, however, detect such an excess between the modern LC line individual and the second Caucasian wisent (Cc2), relative to the modern L line individuals. We can therefore infer that the last surviving Caucasian wisent, KAUKASUS, whose living descendants comprise the modern LC line, was more closely related to Caucasian wisent individual Cc1 that to individual Cc2.

We further investigated Caucasian ancestry in the genome of the modern LC line individual by performing phylogenetic analysis of non-overlapping 1MB genomic blocks. This analysis involved the modern LC line wisent (MdLC), the founding wisent (PLATEN) that was found to be least admixed with Caucasian wisent, Caucasian wisent (Cc1) and domestic cattle, with water buffalo as outgroup. These datasets were mapped to the reference genome of the zebu, *Bos indicus* (Canavez et al., 2012), to benefit from the chromosome-level assembly of that genome. Of the investigated genomic blocks, we find that 22% return the modern LC line and Caucasian wisent as monophyletic (Fig. 6), and may therefore represent introgressed segments of Caucasian wisent ancestry in this modern LC line individual. Around 8% of these blocks are likely to result from incomplete lineage sorting, based on the frequency of occurrence of the opposing topology (Fig. 6), producing an overall estimate of 14% of the genome of the modern LC individual that results from Caucasian wisent admixture, most likely inherited from the bull KAUKASUS. In addition to providing an estimate of admixture proportions, our method is also able to accurately map admixed segments of the genome (Fig. 6a). Many of these segments span multiple megabase blocks. For example, a contiguous 22MB admixed block is observed on chromosome 4, which may span as much as 33MB under the assumption that intervening blocks with missing data are linked to adjacent ones. Relative admixture proportions also vary among chromosomes in this individual. For example, chromosome 27 almost entirely lacks Caucasian wisent ancestry whereas around 50% of chromosomes 4 and 15 are likely derived from admixture.

**Figure 6.**
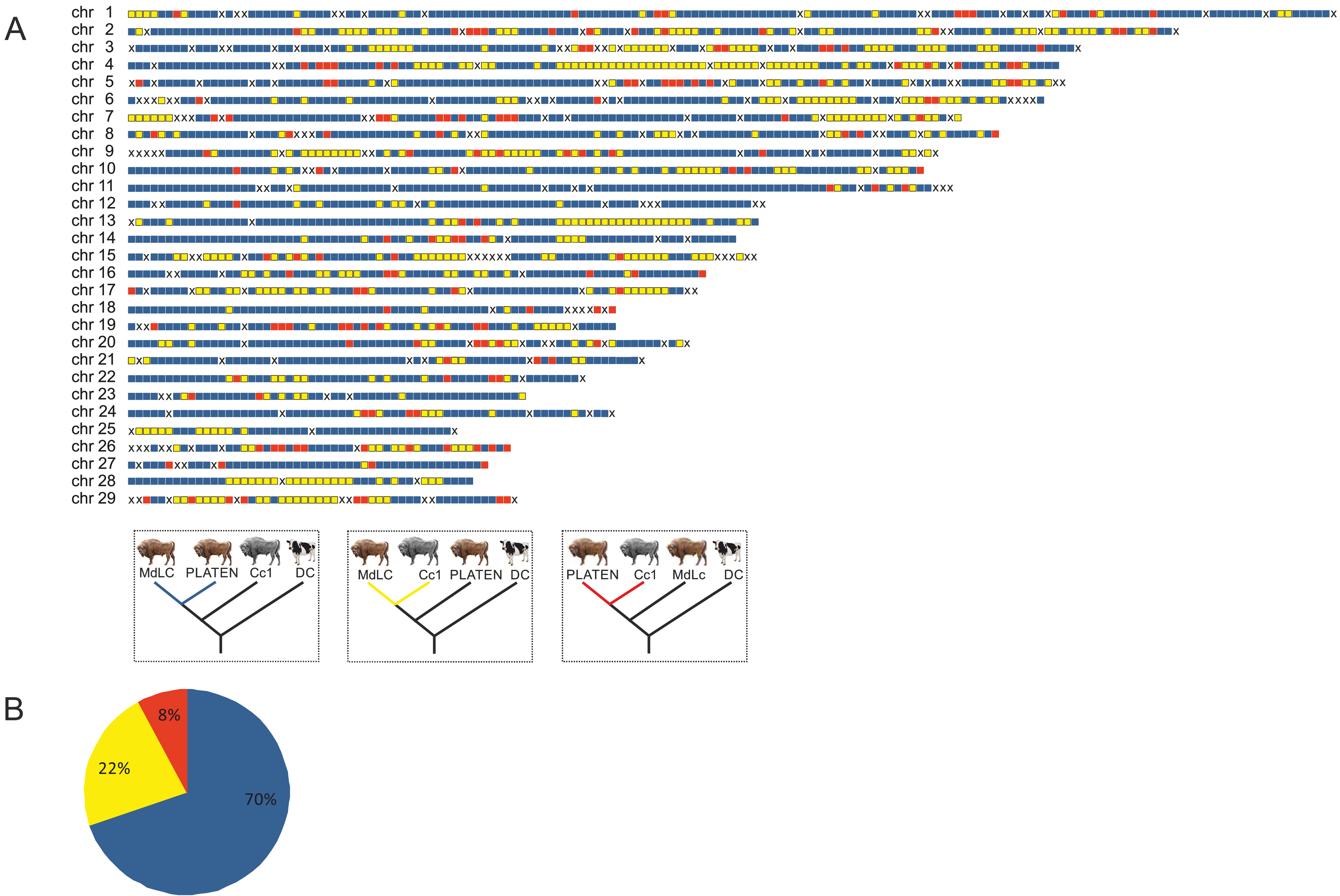
Genomic admixture map (A) of Caucasian bison ancestry in the modern LC line individual (MdLC). Colored blocks indicate 1Mb genomic blocks returning alternative tree topologies, blue blocks are compatible with the species tree; yellow blocks return the monophly of the modern LC line and Caucasian wisent, and likely result from admixture and to a lesser extent incomplete lineage sorting; red blocks return the monophyly of PLATEN and Caucasian wisent and likely result from incomplete lineage sorting. “X” shows blocks with missing data. The pie chart (B) shows the percentage of 1Mb genomic supporting each tree topology identified by colours, according to the key presented above.

Finally, we investigated evidence of Caucasian wisent ancestry in the modern L line. We found that both modern L line wisent share a significant excess of derived alleles with Caucasian wisent (Fig. 5b), relative to founding wisent. Thus, modern L line individuals appear more admixed with Caucasian wisent than the two founding wisent investigated here. This admixture signal could result from either variable admixture proportions among founding individuals, or recent geneflow between the L and LC lines, although *D* statistic comparison of modern individuals failed to detect the latter (see above). We further investigated these alternative hypotheses by comparing the sizes of putatively admixed genomic blocks. Recent geneflow results in large contiguous genomic blocks derived from admixture in the genomes of the recipient population, which are broken up over time as a result of recombination. Pedigree information provides an approximate date for geneflow from Caucasian wisent into the modern LC line around 90 years ago (15-22 generations), the result of which are many intact multi-megabase genomic blocks derived from Caucasian wisent in the modern LC line individual (Fig. 6). We compared the sizes of these blocks with those of putative Caucasian wisent ancestry in a modern L line individual, and found the abundance of large blocks to be considerably lower in the latter (Fig. 7). This rejects recent admixture and instead supports variable admixture proportions among the founding herd in explaining the observed signal of Caucasian wisent admixture in this modern L line individual.

**Figure 7.**
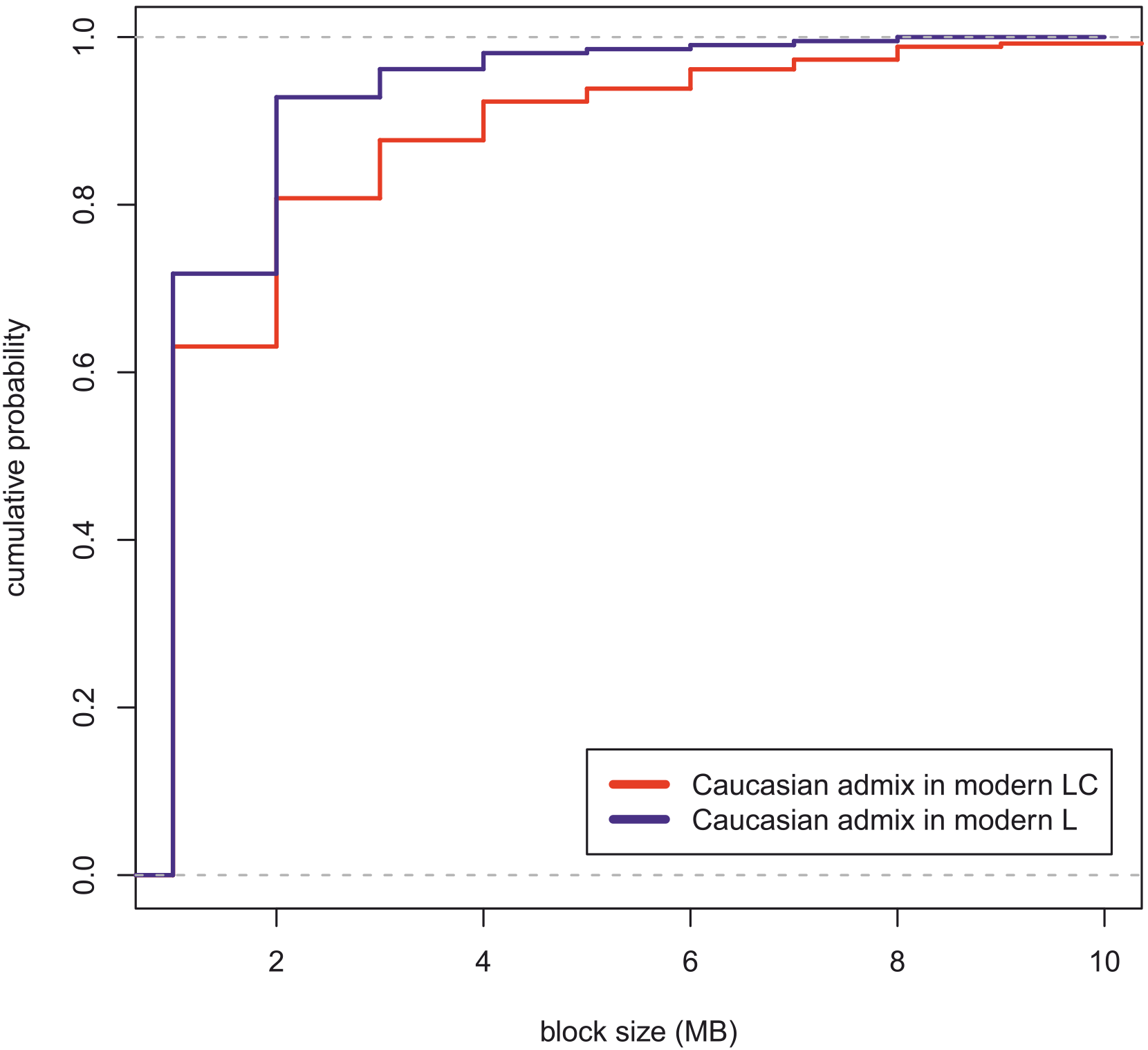
Variation in the sizes of genomic blocks in modern L (blue) and LC (red) likely resulting from Caucasian wisent admixture. Plots show cumulative probability densities calculated at a scale of 1MB. Genomic blocks in the LC line wisent (red) result from admixture occurring around 90 years ago; the lower abundance of larger admixed blocks in the modern L line wisent support that this admixture event preceded the former. The plots have been truncated to aid visualisation, and single blocks of 18MB and 22MB in the LC line individual are not shown. The largest block size detected in the modern L line individual was 8MB.

### Admixture with the cattle/aurochs lineage

We investigated potential admixture between wisent and the cattle/aurochs lineage using pseudo-haploid sequences generated from short read data of two domestic Holstein cows and an ancient aurochs (*Bos primigenius*; Park et al., 2015), the extinct species from which cattle were domesticated and that lived sympatrically with wisent up until its extinction around 400 years ago (van Vuure, 2005). First, we looked for significant differences in derived allele sharing between cattle and wisent, relative to the aurochs. We found that all investigated wisent share a significant excess of derived alleles with cattle relative to aurochs (Fig. 5d). This suggests either admixture between wisent and domestic cattle, or alternatively, admixture with aurochs, if aurochs populations were highly structured and the admixing individuals were from a population more closely related to domestic cattle than the British aurochs used in this analysis.

We also compared derived allele sharing with cattle among the individual wisent. We found that all modern wisent investigated here share a significant excess of derived alleles with cattle relative to any founding or Caucasian wisent (Fig. 5e). Variable admixture was also observed among founding wisent. Specifically, PLANTA shares more derived alleles with domestic cattle than either PLATEN or the Caucasian wisent do. Finally, we estimated the genomic fraction in the modern wisent that could be attributed to cattle/aurochs admixture, above that occurring in the founding wisent, using the 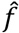 statistic (Durand et al., 2011). This analysis showed that 2.4% to 3.2% of the sampled modern wisent genomes can be attributed to admixture with the cattle/aurochs lineage, respectively, that is not the result of direct inheritance from the sampled founding wisent (Tab. S6).

The increased admixture signal observed in all modern wisent relative to founding wisent could result from either variable cattle/aurochs admixture proportions among the founding herd from which all modern wisent are derived, or alternatively, from recent admixture with modern cattle. However, the fact that we do not find evidence of any complete genomic 1MB blocks resulting from cattle admixture in the modern LC line individual (Fig. 6a) argues strongly against recent cattle admixture, and instead supports variable cattle/aurochs admixture among the founding herd in explaining the excess of derived alleles shared among domestic cattle and modern wisent.

## DISCUSSION

Retracing complex population histories can be challenging. In particular, admixture involving populations or species that are now extinct may be impossible based solely on data from living individuals (Hofreiter et al., 2015). Through the use of low-coverage genomic data from modern and historical wisent, including from the now extinct Caucasian wisent subspecies, we have revealed the complexity of wisent evolution. This complex history involved not only admixture resulting directly from the captive breeding program, but also older processes occurring prior to their extinction in the wild, which included admixture with another bovid lineage (Fig. 8).

**Figure 8.**
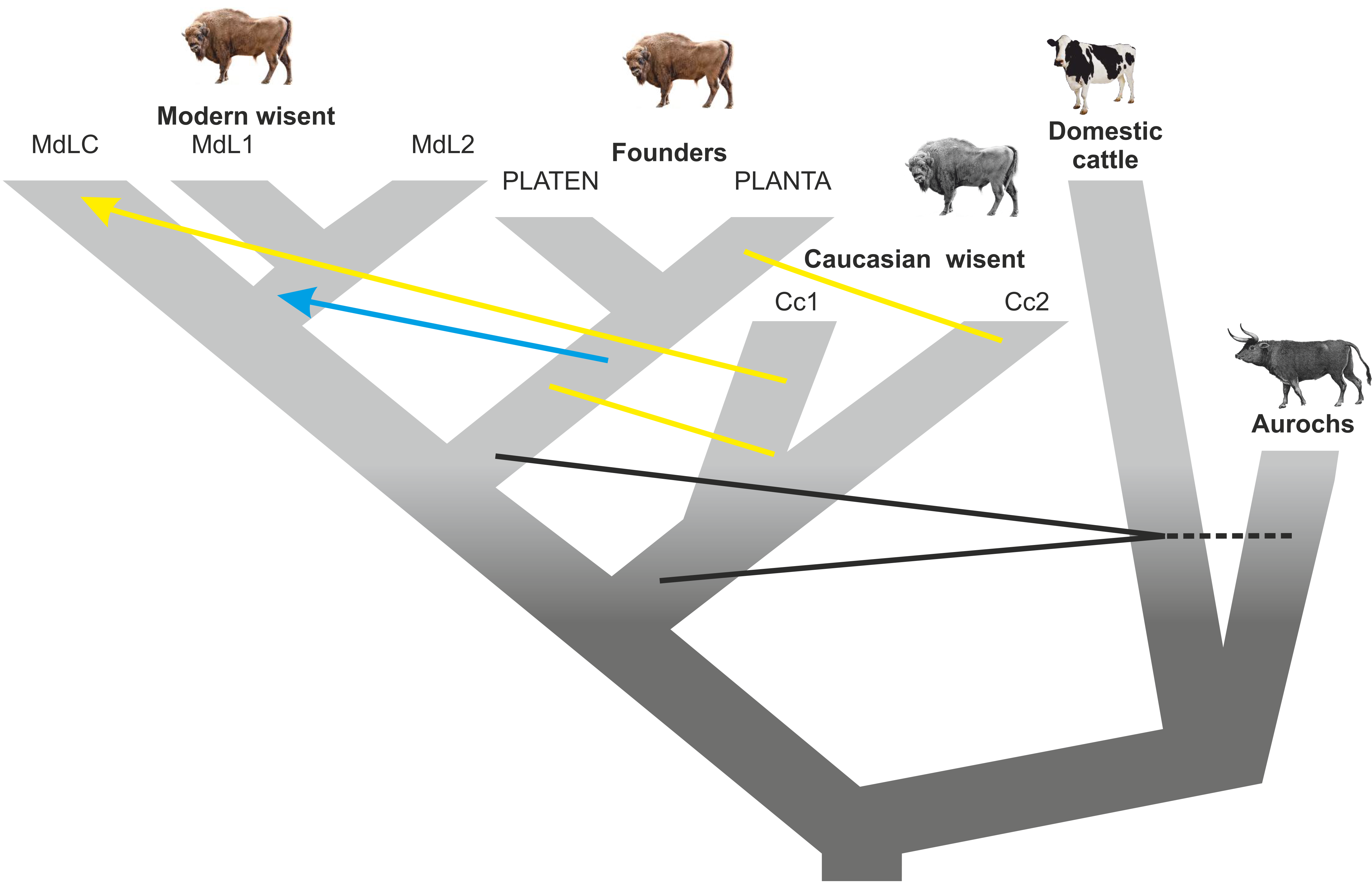
Schematic diagram showing inferred admixture among wisent, and among wisent and the cattle/aurochs lineage. Arrows indicate the direction of the geneflow. Black lines indicate admixture between wisent and cattle/aurochs, yellow lines/arrow – between Caucasian and founding or modern wisent respectively, and the blue arrow – from founders to modern wisent.

### Wisent evolution and admixture

The accepted view of wisent evolution is of two distinct subspecies, Lowland wisent and Caucasian wisent, that both underwent dramatic population declines, with the last few surviving individuals serving as founders of the modern L (Lowland only) and LC (Lowland and Caucasian) lines (Pucek et al., 2004). Our results show that this model is an oversimplification. We find evidence of at least two highly differentiated populations within Caucasian wisent, with one of these showing greater pairwise similarity with Lowland wisent than with the second Caucasian population, at the level of the complete genome (Fig. 2). However, analysis of aligned nuclear genomic blocks from four individuals returns Caucasian and Lowland wisent as reciprocally monophyletic across slightly more than half of the genomic alignment (Fig. 3), providing support that this topology reflects the true history of population divergence. Thus, among any two Caucasian wisent and any two modern (or founding Lowland) wisent, we may expect that any single locus has only around 50% probability of reflecting the true history of population divergence. Moreover, increased sampling of individuals is likely to further reduce this proportion, potentially to such an extent that, at a given sampling level, the true evolutionary history cannot be untangled from the effects of random drift and more recent admixture. This result reinforces the notion that phylogeny-based interpretation may be inappropriate at the level of the complete genome, and that alternative, more flexible models will be required to keep pace with our ability to generate such data (Hofreiter et al., 2015).

A further implied assumption of the traditional view of wisent evolution is that, with the exception of the Caucasian bull KAUKASUS, all founding wisent represented “pure” Lowland wisent (Pucek et al., 2004; Tokarska et al., 2015). On this basis, the modern L line that is derived only from the latter can also be considered as pure Lowland wisent, referable to the subspecies *B. b. bonasus*. Our results also demonstrate that the notion of wisent subspecies purity is flawed in the sense that founding Lowland individuals were in fact admixed with Caucasian wisent to varying degrees. We demonstrate this both directly for the founder PLANTA in comparison to another founder, her son PLATEN, and also indirectly, by the elevated signal of Caucasian wisent admixture in modern L line wisent, relative to these founders, most likely a result of inheritance from other founding individuals not included in this analysis (Fig. 7). The notion of subspecies purity therefore disregards the fact that admixture between Caucasian and Lowland wisent almost certainly occurred prior to the extinction of wisent in the wild, and such admixture could therefore be regarded as part of the normal population processes and dynamics of this species.

The notion of subspecies purity has driven efforts to ensure that free-living L and LC herds do not come into contact (Pucek et al., 2004), and also motivated genetic investigations of living populations that may have been recipients of geneflow from the opposing genetic line (Tokarska et al., 2015). The latter study investigated the modern L line population that currently inhabits the eastern, Belarussian part of the Białowieża Forest. Some individuals were found to possess a microsatellite allele that was common among Caucasian wisent but absent in all studied Lowland wisent, which these authors interpreted as evidence of recent admixture with the modern LC line. Although the individual from this population (modern L line, MdL2) that we sequenced does not possess this putative Caucasian wisent allele, we nevertheless detected evidence of Caucasian wisent admixture above that occurring in founding wisent for this individual (Fig. 5). However, the small size of admixed blocks, in addition to non-significant *D* comparisons of modern lines, supports variable Caucasian wisent admixture among founding wisent in explaining this result, which may also account for the occurrence of putative Caucasian wisent alleles in other individuals from this population. Future studies of such individuals using the methodology applied here would provide a robust test of these alternative hypotheses.

### Wisent conservation and de-extinction

The issue of low genetic variability among living wisent is considered as cause for concern, and has been demonstrated by several population-level studies using various molecular and biochemical markers, such as blood-group systems and blood serum proteins (Sipko et al., 1995; Gębczyński & Tomaszewska-Guszkiewicz 1987; Hartl & Pucek, 1994; Sipko et al., 1997), mtDNA (this study; Tiedemann et al., 1998; Burzyńska et al., 1999; Wójcik et al., 2009; Hassanin et al., 2012), nuclear gene sequences (Sipko et al., 1994; Udina et al., 1994; Kamiński & Zabolewicz 1997; Udina & Shakhaev 1998; Burzyńska & Topczewski, 1999; Radwan et al., 2007; Babik et al., 2012; Hassanin et al., 2013), microsatellites (Gralak et al., 2004; Luenser et al., 2005; Tokarska et al., 2009a; 2009b) and SNPs (Tokarska et al., 2009b, 2015). In apparent contrast to these results, we find relatively high levels of genomic divergence among both modern and founding wisent, being approximately equal to or exceeding that found between pairs of cattle or yak, respectively. Pairwise divergence between modern and Caucasian wisent was found to be even higher, indicating a major loss of genetic diversity from the species as a whole following the extinction of the latter population.

The discrepancy between estimates of genetic variation obtained from population-level studies and from pairwise nuclear genomic divergences, as reported here, may result from several factors. First, application of microsatellites or SNP markers developed for cattle to wisent will likely lead to ascertainment bias (Albrechtsen et al., 2010), particularly if diversity estimates for wisent are then compared back to those obtained for cattle using the same marker set. Admixture with the cattle/aurochs lineage detected in this study adds a further layer of complexity in interpreting data from molecular markers developed for cattle. Second, although genetic diversity and pairwise genetic divergence are both measures of genetic variation in the broad sense, these measures are different and not necessarily interchangeable. Thus, wisent may lack allelic diversity, but the alleles that do exist may be highly divergent from one another. Indeed, the bimodal distributions observed for pairwise comparisons based on transitions and transversions are compatible with high levels of relatedness -- likely inbreeding -- among modern wisent, and low mitochondrial haplotype diversity among modern and historical individuals in comparison to that found among middle Holocene wisent (Fig. S1) provides further evidence of a loss of allelic variation. Under this scenario of low allelic diversity but high nuclear allelic divergence, given continued population expansion and sufficient time, nuclear genomic diversity may potentially be restored as recombination dissects and shuffles divergent chromosomes over successive generations.

The Caucasian wisent is extinct in the wild, but a fraction of its genepool survives in the genomes of modern wisent. Our results provide not only a direct measure of this admixture in a modern LC line individual, but also allow us to map with relative accuracy chromosomal segments that are inherited from Caucasian wisent. Although sometimes controversial, the concept of de-extinction has generated considerable interest (Sherkow & Greely, 2013), and attempts are currently underway to generate animals that, at least superficially, resemble the quagga (*Equus quagga*; Heywood, 2013), an extinct subspecies of plains zebra, and also the aurochs (van Vuure, 2005), by careful selective breeding of their living relatives. In both of these cases, selective breeding and the ultimate success of the project are based solely on morphological criteria. Our study demonstrates that, at least in principle, by generating chromosomal admixture maps for multiple living representatives of the LC line, it would be possible to selectively breed an animal that is, at the genomic level, highly similar to a Caucasian wisent.

### Admixture with the cattle/aurochs lineage

Hybridisation of wild species with their domesticated close relatives is a subject of considerable discussion and concern for conservation management (Ellstrand et al., 2010). Frequently, such events are deemed to be detrimental to the recipient wild species, as is the case for American bison (*Bison bison*) herds found to be admixed with cattle (Halbert & Derr, 2007). However, the introgression of alleles from domestics into wild populations may also provide the basis for adaptation (Fuelner et al., 2013; Anderson et al., 2009), and allow populations to take advantage of new ecological niches (Monzón et al., 2014). In any case, the identification of admixture is the essential first step to guide conservation policy, and measuring admixture proportions among individuals is likely the second essential step for implementing conservation measures. Our results achieve both of these objectives.

We detect a significant proportion of admixture with cattle relative to aurochs in the genomes of all wisent except founder PLATEN. However, since D statistic is a relative test, and PLATEN is the offspring of admixed founder PLANTA, it is reasonable to infer the PLATEN is also admixed to some extent. Although our comparisons consistently found an excess of derived alleles shared between cattle and wisent, relative to aurochs and wisent, it is difficult to completely exclude admixture with aurochs as an alternative explanation, given the lack of knowledge of population structure in aurochs. Testing these two alternatives would require data from additional aurochs populations from within the core distribution of wisent, and would be a valuable direction for future research.

The timing of admixture also has implications for conservation management. Specifically, the removal of individuals resulting from very recent hybridisation may be deemed appropriate (Halbert & Derr, 2007). The small size of cattle admixed blocks in modern wisent (at least undetectable at a 1MB scale) clearly rejects recent cattle admixture for the individuals investigated here. Instead, admixture must have occurred prior to the establishment of the captive breeding program, and the admixture signal detected in modern wisent results from inheritance from the founders that were admixed with cattle to varying degrees. Thus, based on the current evidence, cattle introgression appears of low concern for wisent conservation for the following reasons: 1. admixture does not appear to have occurred since the establishment of the captive breeding program, although screening of additional individuals may be desirable to further support this generalisation; 2. the number of intervening generations separating living wisent from the F1 hybrids is likely sufficient that all living wisent are admixed to some extent (Chang, 1999); and 3. our results may in fact reflect admixture with aurochs, rather than domestic cattle, although this hypothesis requires further investigation.

### Conclusions

The ability to detect admixture is of key importance for both evolutionary and applied conservation studies. However, interpretation of a significant signal of admixture, in terms of both evolutionary inference and the formulation of management strategies, may require information its timing. Using seven low-coverage wisent genomes from both modern and historical wisent we have revealed multiple instances of admixture, but moreover, because the approximate age of introgression of Caucasian wisent into the modern LC line is known, through comparisons of the sizes of likely admixed genomic segments we have inferred the relative ages of other admixture events. This unique historical information, coupled with the ability to recover genomic data from historical samples, establish wisent as an exemplary taxon for the study of admixture in wild populations. As new analytical methods for studying admixture are developed, wisent can serve as a valuable empirical test of both their performance and utility.

## MATERIALS AND METHODS

Complete details of all samples and specimens used in this study are shown in Table 1.

### Laboratory methods, modern samples

DNA was extracted from tissue samples of three modern wisent using either a DNeasy Blood & Tissue Kit (Qiagen) according to the manufacturer’s instructions (sample MdL1) or by phenol/chloroform extraction (Sambrook & Russell, 2001). We mechanically sheared the DNA of the modern samples using a Covaris S220 sonicator to an average fragment length of 500 bp and prepared indexed Illumina libraries from 500 ng of each modern DNA extract using a published double-stranded protocol (Meyer & Kircher, 2010) with modifications (Fortes & Paijmans, 2015). Library molecules from 450 bp to 1000 bp were then selected using a Pippin Prep Instrument (Sage Science).

### Laboratory methods, historical samples

DNA extraction from four museum specimens as well as sequencing library preparation steps preceding amplification were performed in a dedicated ancient DNA laboratory (Evolutionary Adaptive Genomics Group, Potsdam University, Germany). DNA extracts were prepared from horn and bone powder obtained by grinding in a mixer mill (MM 400, RETSCH). DNA extraction followed the protocol of Dabney et al. (2013a), except for horn samples where we used a different digestion buffer containing 10mM Tris buffer (pH 8.0), 10 mM NaCl, 5 mM CaCl_2_, 2.5 mM EDTA (pH 8.0), 2% SDS (Shapiro & Hofreiter, 2012). The museum samples were already fragmented due to degradation, so were not sonicated. We used 25 μl of each DNA extract to construct single-stranded indexed Illumina libraries according to the protocol of Gansauge & Meyer (2013).

### Sequencing

Final library concentrations and the distribution of insert sizes were determined using a 2200 TapeStation (Agilent Technologies) and Qubit HS-assay (Thermo Fisher Scientific), respectively. Each library was then sequenced using an Illumina NextSeq 500 instrument. For modern libraries we used a High Output Kit (75 bp paired-end sequencing), for libraries obtained from historical horn samples we used High Output Kits (75 bp single-end and 150 bp paired-end) and each library built from historical bone samples was sequenced separately with High Output Kits (75 bp single-end and paired-end). Full details of sequencing results are provided in Supporting Tables S2 and S3.

### Data processing, mapping and pseudo-haploidisation

For paired-end data, we trimmed adapter sequences and merged overlapping read pairs using SeqPrep (https://github.com/jstjohn/SeqPrep), requiring a minimum read length of 30 bp (-L 30), minimum overlap of 15 bp (-o 15), and a minimum merge quality of 13 (-q 13). Adapters occurring at the 3’ ends of single-end reads were trimmed using cutadapt (Martin, 2011), also requiring a minimum length of reads of 30 (-m 30). We then mapped the resulting data to the zebu (*Bos indicus*; GenBank accession no. GCA_000247795.2) and water buffalo (*Bubalus bubalis*; GenBank accession no. GCA_000471725.1) nuclear genomes and wisent mitochondrial genome (KW, unpublished) using BWA aln version 0.7.8 (Li & Durbin, 2009) with default 0.04 mismatch value. We removed duplicate reads likely resulting from PCR amplification using samtools rmdup (Li et al., 2009). Detailed descriptions of the mapping results are provided in Supporting Tables S2 and S3. We then generated pseudo-haploid sequences as described by Cahill et al. (2015) and used these for further analysis.

### Pairwise genomic divergence

Pairwise genomic divergence was calculated by dividing genomic alignments into nonoverlapping 1MB blocks and calculating the proportion of transversions, or transitions plus transversions (comparisons of modern individuals only), for each pair of individuals, accounting for the presence of missing data. Blocks with > 75% missing data were disregarded. Probability densities were generated by kernel density estimation in R (R Core Team, 2014) using default parameters. Full details of comparative data generated for domestic cattle and yak (data from the NCBI Short Read Archive) are provided in Supporting Table S4.

### Mitochondrial genome analysis

In addition to the mitochondrial genomes generated from the seven modern and four historical specimens, we generated four mitogenomes from ancient middle Holocene wisent individuals (Tab. 1). These ancient samples were radiocarbon dated at either the Oxford University Radiocarbon Accelerator Unit (Oxford, UK) using ultrafiltered collagen and accelerator mass spectrometry (Ramsey et al. 2004a, 2004b) or the VERA-Laboratorium Institut fur Isotopenforschung und Kernphysik (Vienna, Austria). We calibrated radiocarbon dates using the IntCal13 calibration curve (Reimer et al., 2013) in OxCal v4.2 (https://c14.arch.ox.ac.uk/oxcal/OxCal.html).

DNA was extracted from the Holocene *B. bonasus* samples (Bb1-Bb3) at the Henry Wellcome Ancient Biomolecules Centre (Oxford University, UK), following Shapiro et al. (2004). We extracted the Holocene *B. b. caucasicus* sample (Cc3) in the specialist Paleogenomics facility at UC Santa Cruz, following Rohland et al. (2010). DNA library construction, mitochondrial target enrichment, sequencing, and sequence data processing protocols for the four Holocene samples followed approach four in Heintzman et al. (2016), except that the whole mitochondrial genome consensus sequence was retained. The mean read depth of these Holocene consensus sequences ranged from 14.1 to 165.7x. The consensus sequences for the four Holocene and one historic Caucasian bison have been deposited in GenBank with accession numbers XXXXXXXXX-XXXXXXXX. The mitogenomic sequences from the remaining modern and historic samples were identical to either haplotypes previously published or the historic Caucasian bison haplotype (Tab. S1).

We assessed phylogenetic relationships among wisent mitochondrial haplotypes, as well as their placement within the wider bovin (tribe Bovini) tree. Wisent sequences were aligned with those of 12 other bovin taxa, including the extinct steppe bison (*Bison priscus*) and aurochs (*Bos primigenius*) (Tab. S1). We excluded two previously published wisent mitochondrial genomes from the phylogenetic analysis (GenBank accessions: HQ223450, HM045017/NC_014044), as these sequences were considered problematic. Specifically, HQ223450 has multiple insertions totaling 9 bp in the ND4 coding region, and HM045017/NC_014044 has multiple indels and point mutations concentrated in the large rRNA and ND3 coding regions. Sequence alignment, partitioning, model testing, and phylogenetic and associated statistical support methods followed the ordinal-level analyses of Heintzman et al. (2015), except that we used the *B. bison* reference mitochondrial genome (NC_012346) for partitioning. We selected the following models of molecular evolution for the six partitions: GTR+I+G (CP1, 3803 bp; rRNAs, 2541 bp), GTR+G (CP3, 3803 bp), HKY+I+G (CP2, 3803 bp; tRNAs, 1526 bp; control region, 927 bp). We used the saola (Pseudoryx nghetinhensis; Tab. S1) as outgroup in both the maximum likelihood and Bayesian analyses, following Bibi (2013).

### D statistic tests

The *D* statistic involves 4 genomes: a genome from each of two sister populations (P1 and P2), a genome from a third population as the potential source of introgression (P3), and an outgroup genome (O) to identify the ancestral state (identified as the A allele). We identified variable positions at which P3 possessed the derived allele (B) and presence of the derived allele is variable among P1 and P2, leading to two possible patterns: either ABBA or BABA. Under the scenario of incomplete lineage sorting without geneflow these patterns should occur with equal frequency and the expected *D* value will be zero. An excess of ABBA or BABA patterns is interpreted as evidence of admixture. However, it might also arise from nonrandom mating in the ancestral population due to population structure (Eriksson & Manica, 2012). To determine the ancestral state we used the water buffalo genome. In all tests involving data mapped to the zebu genome, we took into consideration the autosomes only. We performed a total of 105 comparisons considering all possible combinations of wisent, all wisent with either domestic cattle or aurochs as candidate admixer, and domestic cattle and aurochs with all wisent as candidate admixer. These results are reported in Supporting Table S4. 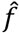 test (Green et al., 2010; Durand et al., 2011) was used to estimate the proportion of the genome derived from admixture. This test requires two individuals of the candidate introgressor species that are not themselves admixed. For our datasets this was possible only for admixture involving the cattle/aurochs lineage. For both *D* statistic and 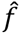 test, significance was assessed using a weighted block jackknife using 1Mb blocks (Green et al., 2010; Durand et al., 2011). The weighted block jackknife tests if admixture signals are uniform across the whole genome and therefore reflect the same population history. By removing one at a time blocks of adjacent sites (larger than the extent of linkage disequilibrium) and computing the variance of the *D* statistic or 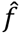 values over the entire genome *M* times leaving each block of the genome in turn, and then multiplying by *M* and taking the square root we generated the standard error. The number of standard errors by which *D* or 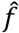 differs from zero is the *Z* score. The results with *Z*-scores greater than 3 in absolute value were qualified as statistically significant (Green et al., 2010).

### Nuclear genome phylogenetic tests

The aligned pseudohaploid sequences were divided into non-overlapping blocks of 1MB. If each of the five taxa contained no more than 50% gaps within a window, the sequence data were recoded into binary characters to only score transversions (Rs: 0, Ys: 1), otherwise the window was recorded as having insufficient data. A Maximum Likelihood phylogeny under the BINGAMMA model and with the water buffalo as outgroup was then computed for each alignment with sufficient data using RaxML (Stamatakis, 2014). The topology of each phylogeny was evaluated using a custom Perl script that made use of the ETE3 software (Huerta-Cepas et al., 2016). The lengths of admixed genomic regions was estimated by counting the number of consecutive 1MB blocks returning the respective tree topology. Due to the presence of blocks with insufficient data, these measurements are likely to be underestimates. Evaluation of the lengths of genomic regions was conducted using the empirical cumulative distribution function in R, with default parameters.

## ACKNOWLEDGEMENTS

We would like to thank Andrzej Imiołczyk and his colleagues from the Specimen Preparation Laboratory of the Upper Silesian Museum in Bytom. We thank Andrew Fields and Joshua Kapp for technical assistance. The study has been funded by National Science Center (NCN) grant 2011/03/N/NZ8/00086 to KW.

## AUTHOR CONTRIBUTIONS

Conceived the study – KW, JMS, AB, MH

Designed laboratory experiments – AB, KW, JLAP, PDH

Performed lab work – KW, UT, GX, PDH, BS

Coordinated data analysis – KW, SH, AB, JAC, JLAP, PDH

Performed data analysis – KW, SH, AB, PDH

Coordinated writing of the manuscript – KW, AB, MH

Obtained funding – KW, JMS, MH

Provided samples – GB, ANB, JJC, RD, NM, HO, MT, STT, JMW, WŻ

All authors read, gave comments and helped revise the final version of the manuscript.

## SUPPORTING INFORMATION

Figure S1. A phylogeny of wisent (A) and the Bovini (B), inferred from a partitioned maximum likelihood (ML) analysis of whole mitochondrial genomes. (A) is an expansion of the region in the blue box in (B). The green box in (A) highlights the two haplotypes found in sampled historic and modern individuals: one in Caucasian wisent (Cc2) and the other one identical to haplotype published already (JN632602; Hassanin et al., 2012) in all remaining historical and modern wisent (see Table S1). Tips are coloured based on whether the haplotype occurs in modern individuals (black), or only in historical or ancient individuals, and therefore likely extinct (red). The outgroup, Pseudoryx nghetinhensis, is not shown. Branch support is indicated by bootstrap percentages based on 500 ML bootstrap replicates (above branches) and Bayesian posterior probabilities (below branches).

Figure S2. DNA fragmentation (upper four plots) and deamination (lower two plots) patterns of founding wisent, PLANTA, sequencing reads. For deamination plots, red lines show the frequency of C to T substitutions (Y axes) in the sequenced historical DNA fragments relative to the reference genome at the 5’ (left plot) and 3’ (right plot) fragment ends. X axes show sequenced positions moving internally from the 5’ (positive values) and 3’ (negative values) fragment ends. Elevated rate of C to T substitutions at fragment ends are indicative of DNA damage.

Figure S3. DNA fragmentation and deamination patterns of founding wisent, PLATEN, sequencing reads.

Figure S4. DNA fragmentation and deamination patterns of Caucasian wisent, Cc1, sequencing reads.

Figure S5. DNA fragmentation and deamination patterns of Caucasian wisent, Cc2, sequencing reads.

Table S1. Species and accession data for sequences included in the mitochondrial genome phylogenetic analysis.

Table S2. Full details of sequencing and mapping results for modern wisent samples. SR – merged reads, PE – unmerged reads.

Table S3. Full details of sequencing and mapping results for archival wisent samples. SR – merged reads, PE – unmerged reads.

Table S4. Full details of mapping results for data downloaded from the NCBI Short Read Archive. SR – merged reads, PE – unmerged reads.

Table S5. Detailed *D* statistic results. In red significant results are shown. Results highlighted in grey are presented also in Figure 5.

Table S6. 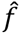 results. In the last column percentage of the genome resulted from hybridisation is given.

